# Predicting the infecting dengue serotype from antibody titre data using machine learning

**DOI:** 10.1101/2024.05.23.595461

**Authors:** Bethan Cracknell Daniels, Darunee Buddhari, Taweewun Hunsawong, Sopon Iamsirithaworn, Aaron R Farmer, Derek A.T. Cummings, Kathryn B. Anderson, Ilaria Dorigatti

## Abstract

The development of a safe and efficacious vaccine that provides immunity against all four dengue virus serotypes is a priority, and a significant challenge for vaccine development has been defining and measuring serotype-specific outcomes and correlates of protection. The plaque reduction neutralisation test (PRNT) is the gold standard assay for measuring serotype-specific antibodies, but this test cannot differentiate homotypic and heterotypic antibodies and characterising the infection history is challenging. To address this, we present an analysis of pre- and post-infection antibody titres measured using the PRNT, collected from a prospective cohort of Thai children. We applied four machine learning classifiers and multinomial logistic regression to the titre data to predict the infecting serotype. The models were validated against the true infecting serotype, identified using RT-PCR. Model performance was calculated using 100 bootstrap samples of the train and out-of-sample test sets. Our analysis showed that, on average, the greatest change in titre was against the infecting serotype. However, in 53.4% (109/204) of the subjects, the highest titre change did not correspond to the infecting serotype, including in 34.3% (12/35) of dengue-naïve individuals. The highest post-infection titres of seropositive cases were more likely to match the serotype of the highest pre-infection titre than the infecting serotype, consistent with original antigenic sin. Despite these challenges, the best performing machine learning algorithm achieved 76.3% (95% CI 57.9-89.5%) accuracy on the out-of-sample test set in predicting the infecting serotype from PRNT data. Incorporating additional spatiotemporal data improved accuracy to 80.6% (95% CI 63.2-94.7%), while using only post-infection titres as predictor variables yielded an accuracy of 71.7% (95% CI 57.9-84.2%). These results show that machine learning classifiers can be used to overcome challenges in interpreting PRNT titres, making them useful tools in investigating dengue immune dynamics, infection history and identifying serotype-specific correlates of protection, which in turn can support the evaluation of clinical trial endpoints and vaccine development.

## Introduction

Dengue is an arboviral infection that has expanded globally in the last 50 years, with an estimated 105 million cases annually (95% confidence interval (CI) 95-114) (1). Despite this, there are currently no specific antiviral treatments or vaccines in widespread use. Dengue is caused by four antigenically distinct virus serotypes (DENV-1-4), which interact immunologically. Infection results in protective and durable homotypic immunity, although homotypic reinfections may occur (2,3). Conversely, heterotypic immunity following a primary infection is temporary, and secondary infections are associated with the potential for disease enhancement due to antibody dependent enhancement (4) which increases viral replication (5), and original antigenic sin (6) which skews secondary immune responses towards the primary infecting serotype. Despite the higher likelihood of disease, a secondary infection also induces a broadly neutralising heterotypic response, to the extent that tertiary and quaternary infections are rarely severe (7).

To avoid enhancement, both the dengue vaccines licenced to date, Dengvaxia and Qdenga, aim to induce balanced immunity against all four serotypes. However, achieving and measuring this has proven challenging. While homotypic antibodies correlate with protection (2), heterotypic antibodies may be broadly protective (following the secondary infection) or enhance disease (following the primary infection), due to differences in their avidity, antigenic targets, and concentration (8,9). The plaque reduction neutralisation test (PRNT), the gold standard for neutralising antibody measurement, cannot differentiate between homotypic and heterotypic antibodies. Consequently, immunogenicity endpoints in vaccine trials rely on total serotype-specific neutralising antibody titres and rates of tetravalent seroconversion, measured using the PRNT or similar neutralisation assays. However, these endpoints have limitations; neutralising titres may be associated with enhancement (10,11) and the commonly used seroconversion threshold (PRNT titre >10) does not correlate with disease protection (10).

This challenge was highlighted by the phase III trial of Qdenga. In the third year, tetravalent seropositivity was 81% in baseline seronegatives, suggesting broad protective immunity against all serotypes in most recipients (12). However, vaccine efficacies in year 3 against DENV-1 to -4 were 17%, 85%, 10% and -99%, respectively. A study of the vaccine’s functional response revealed that most of the serotype-specific antibodies targeted DENV-2 (13), suggesting that seronegative protection against the other serotypes might partly arise from a transient heterotypic response against the DENV-2 component of the vaccine. Additionally, the first licensed vaccine, Dengvaxia, is not in widespread use due to the risk of enhanced disease in subjects who were dengue-naïve (seronegative) at the time of vaccination (14). These results emphasise the need to better understand serotype-specific immunity. While novel assays have been developed to measure serotype-specific antibodies (13,15), they are infeasible for large-scale phase III trials. As such, serology remains central to inferring subclinical infections and estimating vaccine efficacy against infection (16), defining vaccine immunogenicity (17), identifying correlates of protection (18), and studying dengue dynamics in population-level studies (19).

We therefore aimed to test the utility of machine learning classifiers in identifying the infecting serotype from PRNT data. The analysis of PRNT antibody titres collected from a prospective cohort of Thai schoolchildren over five years (20) shows that cross-reactive titres, even in seronegative individuals, can complicate the interpretation of serotype-specific titres. Building on work by van Panhuis *et al.* (21), we trained machine learning classifiers to predict the infecting serotype from pre- and post-infection PRNT titres with high accuracy. We compared these approaches with earlier methods based on multinomial logistic regression (21).

## Results

### Study characteristics

204 symptomatic RT-PCR confirmed dengue cases were recorded between 1998 and 2002 (**Supplementary Figure 1**). Of these, 35 cases were in individuals classified as seronegative prior to infection (pre-infection titres < 10 for all 4 DENV serotypes), although only four of these cases were classified as primary cases according to their ratio of IgM/IgG antibodies (**Supplementary Figure 1a**). The remaining 169 cases were in seropositive individuals (**Supplementary Figure 1b**), 48·5% of which had pre-infection titres ≥ 10 to JEV. The age of infected children ranged from 7-15 years, with a mean age of 10 years (**Supplementary Figure 1c and 1d**). All four DENV serotypes were observed, with DENV-2 being the most prevalent (49%), followed by DENV-3 (32%), DENV-1 (17%) and DENV-4 (2%).

### Antibody dynamics

**Fig. 1** shows the average pre- and post-infection log_10_ reciprocal neutralising PRNT titres, stratified by the infecting serotype. On average, the highest change in titre was against the infecting serotype. Consistent with antibody cross-reactivity, there were rises in neutralising titres against all serotypes, with similar post-infection PRNT titre values against DENV-1, -2 and -3, irrespective of the infecting serotype. Of the 35 cases in seronegative individuals, only one, four or nine had monotypic responses, depending on whether a monotypic response was defined as having a PRNT titre ≥ 10 to only one serotype (22), a PRNT titre ≥ 10 to more than one serotype but a PRNT titre ≥ 80 to only one serotype (20), or a PRNT titre ≥ 10 to more than one serotype but a PRNT titre five-fold higher to only one serotype (23).

**Figure 1:**
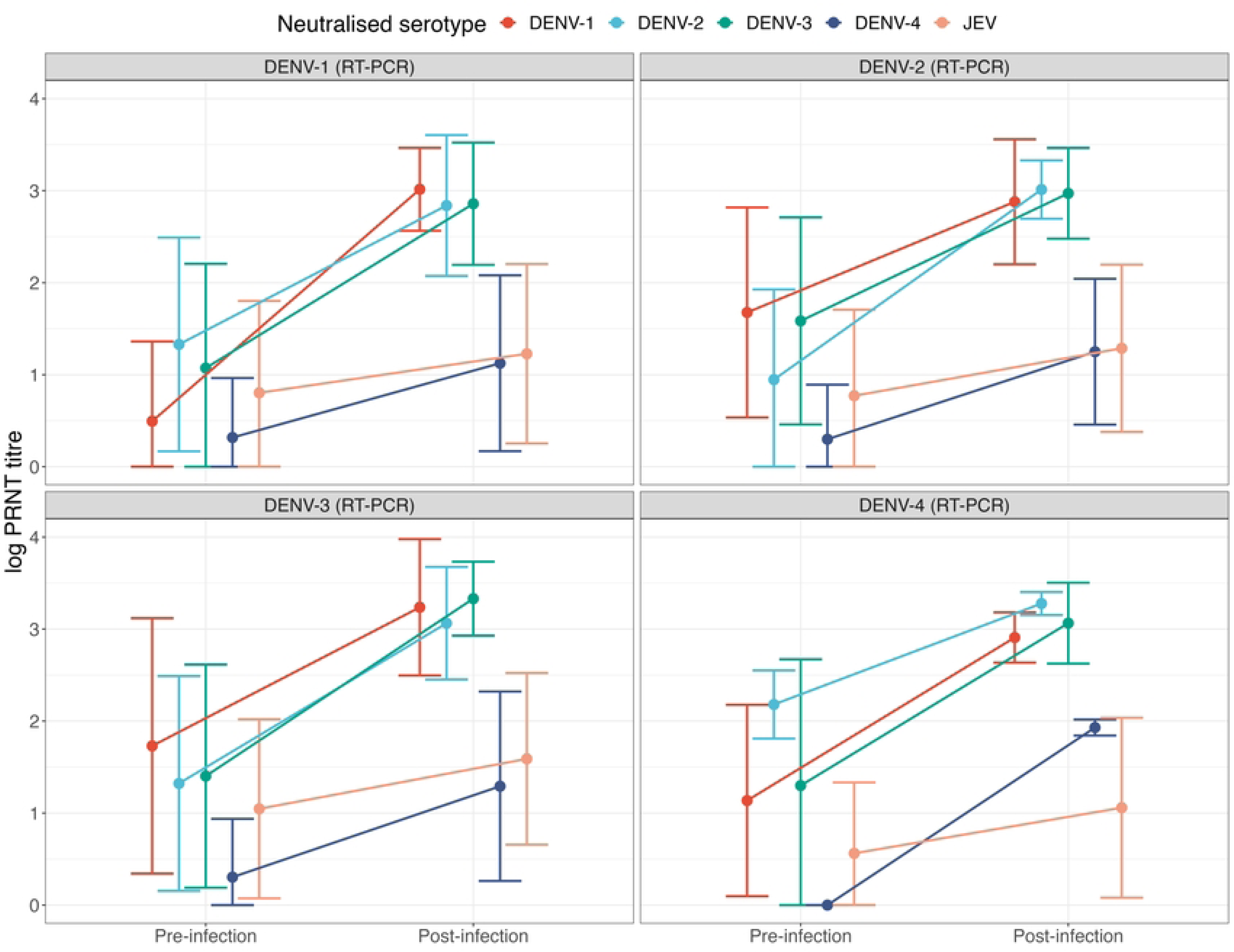
Mean (point) ± standard deviation (error bar) of the pre- and post-infection log PRNT titre, by infecting DENV serotype as quantified by RT-PCR. PRNT: plaque reduction neutralisation test. DENV: dengue virus (DENV1: red, DENV2: blue, DENV3: green, DENV4: dark blue. JEV: Japanese encephalitis virus: orange. RT-PCR: reverse transcriptase polymerase chain reaction.

Among the seropositive cases, 29 (17.2%) had the highest neutralising titre tied between two or more serotypes, due to right-censoring of antibody titres at either 2560 or 10240 (**Fig. 2a**). For the remaining seropositive cases, 76.3% had the highest post-infection titre against a non-infecting serotype, and in 75.5% of these, the serotype with the highest post-infection titre was also the one with the highest pre-infection titre (**Fig. 2a**). Thus, in over half of the seropositive cases without tied highest titres, the highest post-infection titre corresponded to the highest pre-infection titre, rather than to the infecting serotype.

**Figure 2:**
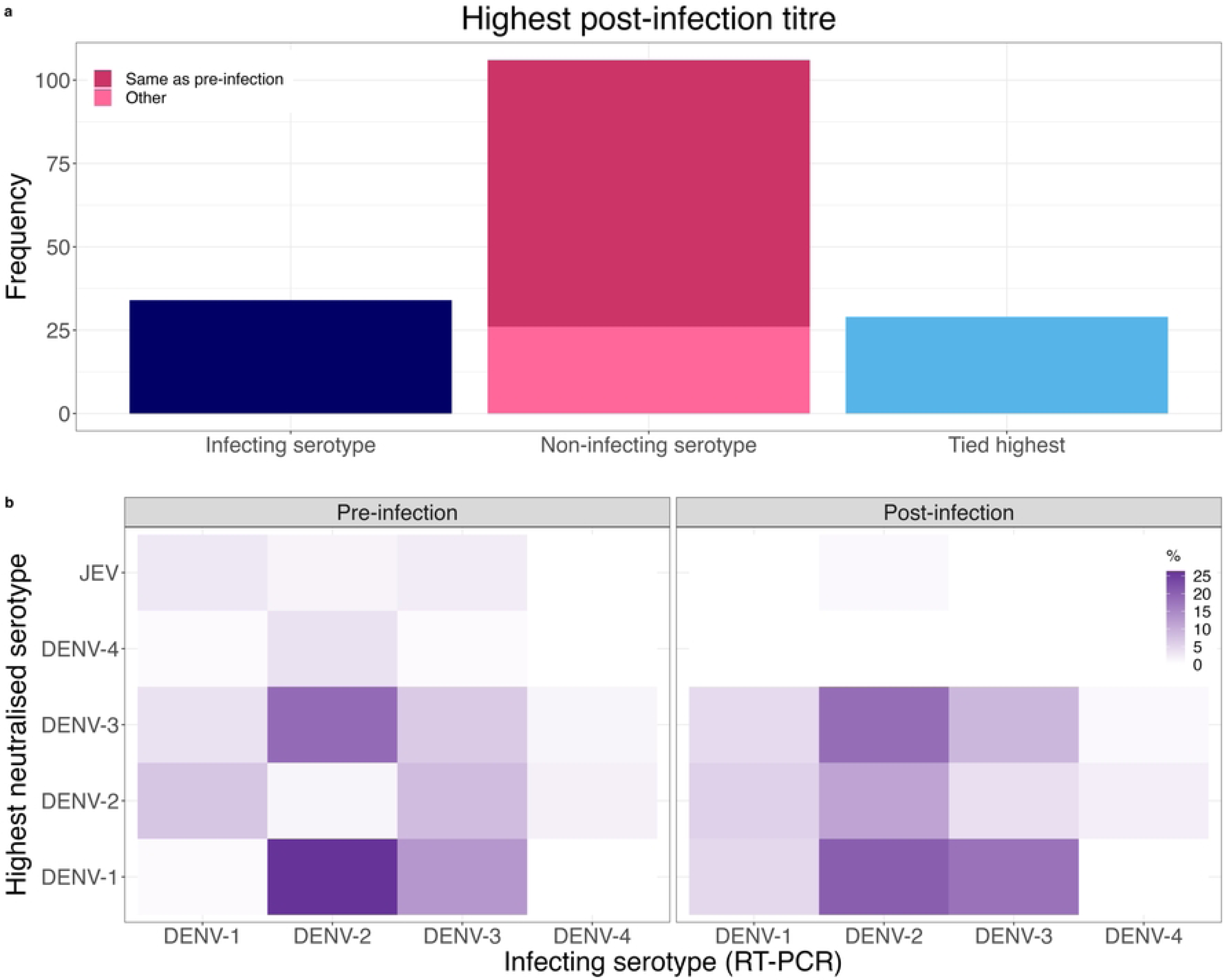
Highest pre- and post-infection log PRNT titres in cases in seropositive individuals (N = 169). **a)** The frequency of subjects whose highest post-infection titre was against the infecting serotype, a non-infecting serotype or whose highest titre was tied between one or more serotypes due to maximum PRNT dilution. **b)** The percentage of subjects whose highest pre- and post-infection titres were against each serotype and JEV, stratified by the infecting serotype. Tied highest titres are split. Seropositive individuals were defined as those with a pre-infection titre ≥ 10 for at least one DENV serotype. PRNT: plaque reduction neutralisation test. RT-PCR: reverse transcriptase polymerase chain reaction. DENV: dengue virus. JEV: Japanese encephalitis virus.

The mean of the log_10_ highest pre-infection reciprocal titre was significantly higher (Student’s T test, p value = 0.01) in cases with a non-infecting serotype as the highest post-infection titre (mean and standard deviation 2.6 and 0.7) compared to those whose highest post-infection titre matched the infecting serotype (mean and standard deviation 2.2, 0.6). **Fig. 2b** shows the frequency of the highest neutralised serotype, pre- and post-infection (cases where the highest titre is against the infecting serotype are along the diagonal). Except for a DENV-3 infection, the highest pre-infection titre was never against the infecting serotype (left hand panel). Notably, a high percentage of DENV-2 infections had highest pre- and post-infection titres against DENV-1 and DENV-3. Additionally, a higher frequency of DENV-3 cases had a highest titre against DENV-1 following the DENV-3 infection than before (**Fig. 2b**).

**Fig. 3** shows the highest PRNT titre change for each case, stratified by the infecting serotype. In 57·4% of seropositive individuals the highest titre changes were against a non-infecting serotype (87/169) or JEV (10/169) (**Fig. 3a**). In seropositive individuals infected with DENV-3 the highest change in titre was equally likely to be against DENV-2 or DENV-3. Surprisingly, even in seronegative cases, the highest titre change did not always correspond with the infecting serotype (**Fig. 3b**). Notably, the highest titre change was against DENV-3 in multiple DENV-1 and DENV-2 infections. Together, these findings suggest that although on average, the greatest change in titre is against the infecting serotype (**Figure 1**), this is frequently not the case, even in individuals who were seronegative prior to infection.

**Figure 3:**
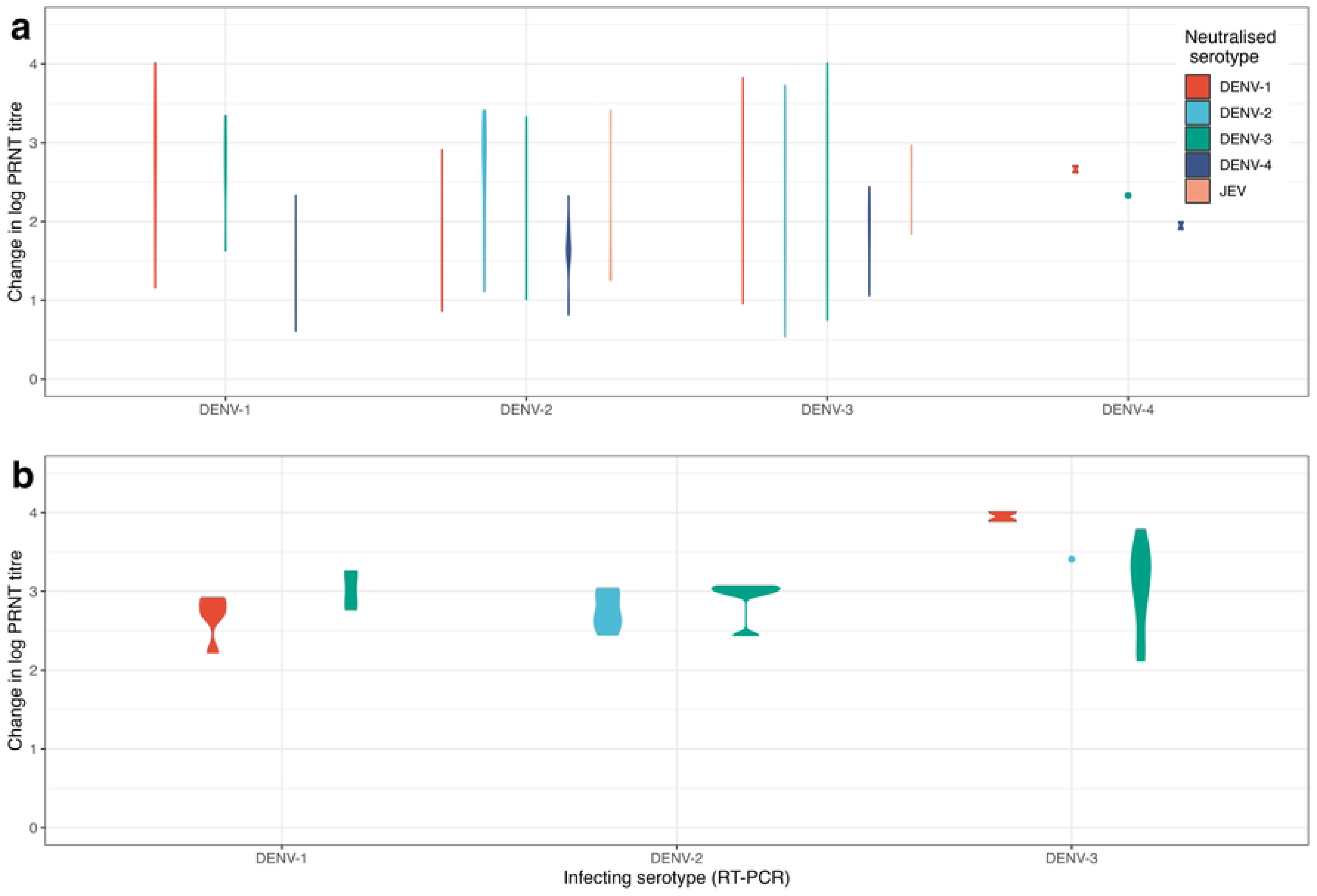
Greatest change in log PRNT titres following infection with a DENV serotype, as quantified by RT-PCR in (a) cases in seropositive individuals and (b) cases in seronegative individuals. Seropositive individuals were defined as those with a pre-infection titre ≥ 10 for at least one DENV serotype. The plot area is proportional to the number of seropositive / seronegative cases. Points indicate a single case. PRNT: plaque reduction neutralisation test. DENV: dengue virus. JEV: Japanese encephalitis virus. RT-PCR: reverse transcriptase polymerase chain reaction.

### Relationship between timing of titre measurement and change in PRNT titre

The time between measurement of pre- and post-infection titres and the date of infection varied within the study, ranging from 9-281 days (with a single outlier of 584 days between measurement of the pre-infection titre and a DENV-3 infection). **S2a Fig.** shows how the average change in log_10_ PRNT titre tends to increase as the number of days between measurement of the pre-infection titre and infection increase, suggesting the decay of antibodies induced against a past infection is associated with a greater increase in antibody titre against the subsequent infection. Conversely, the average change in titre decreased as the number of days between infection and measurement of post-infection titre increased, consistent with decaying titres (**S2b Fig**).

### Predicting the infecting serotype

**Fig. 4** shows an overview of the algorithm used to develop and validate the performance of four machine learning classifiers (random forest RF, gradient boosting machines GBM, artificial neural network ANN, and support vector machines SVM) compared to MLR in predicting the infecting serotype from neutralising antibody titres (including the pre- and post-infection titres, **Fig. 1**, the change in titre, **Fig. 3**, and the number of days between infection and titre measurement, **S2 Fig.)** . **Fig. 5** shows the mean and 95% CI of the performance of each classification model across all four scenarios. In scenario A, which utilised all titre predictor variables (**S1 Table**), the best-performing classifier was GBM, achieving an accuracy of 76.3% (95% CI 57.9-89.5%) on the test set. The kappa statistic was 0.59 (95% CI 0.25-0.83). Training performance was similar (accuracy 74.6% 95% CI 72.7-77.8% and kappa statistic 0.58, 95% CI 0.54-0.64), suggesting that the models are not overfitting to the training data (**Fig. 5**). The accuracy and kappa statistic of MLR on the training set were 65·8% (95% CI 63·0 -68·4%) and 0·46 (95% CI 0·41-0·50), significantly lower than all the machine learning classifiers training set performances. The performance of the machine learning classifiers on the test set was also better than MLR, however this difference was not significant, due to the wide 95% CI of the estimates of model performance (**Fig. 5a**). To investigate the wide 95% CI estimates of model performance on the test set, we fit the GBM model to the scenario A predictors using different train/test proportions (**Fig. 5b**). Increasing the size of the test set from 10% reduced uncertainty in the performance estimates, however, to maximise the data available to train the models given the small size of the dataset, a 90%/10% train and test split was used in the main analysis.

**Figure 4:**
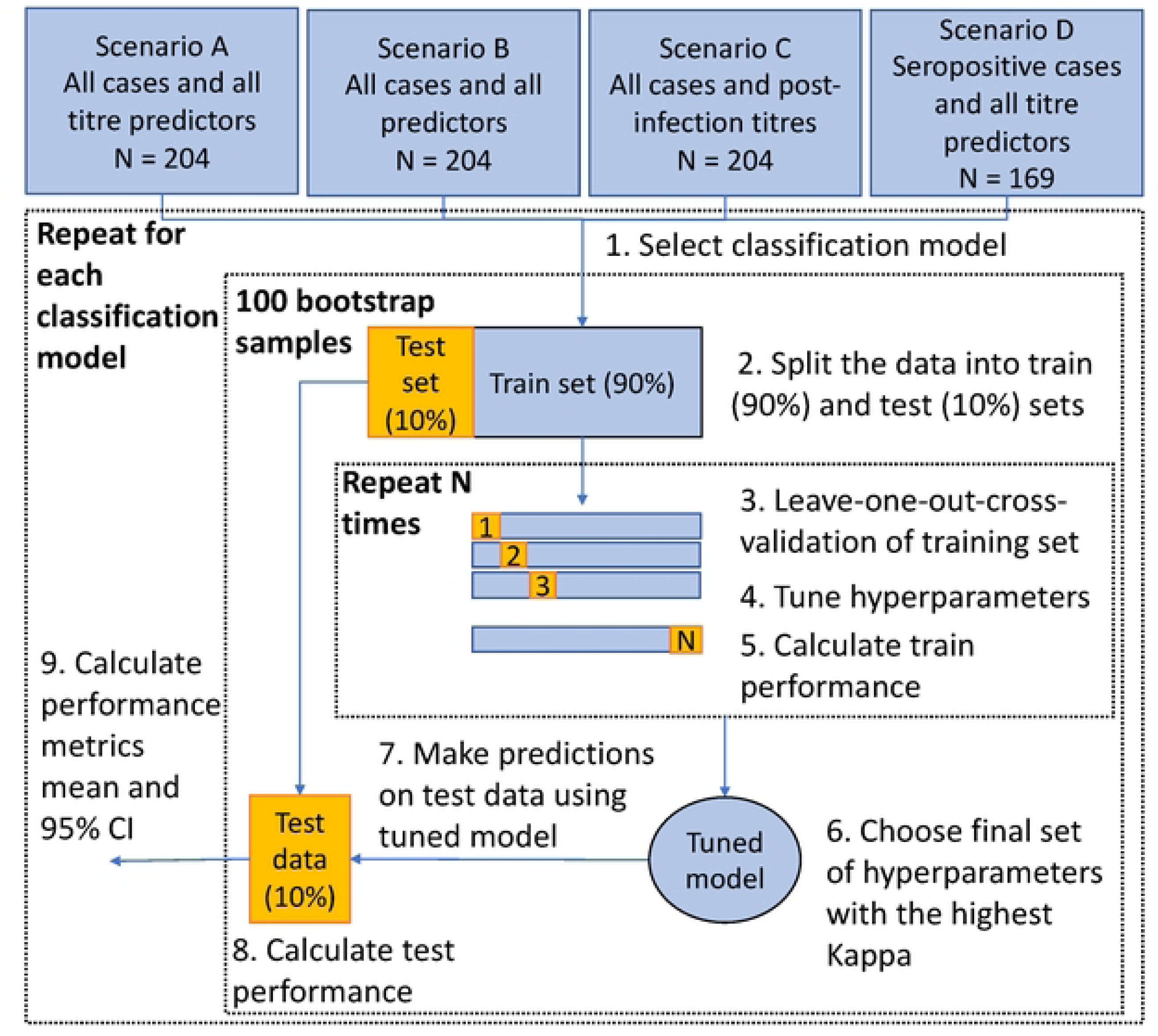
Algorithm of model development and validation. For each scenario and each classifier (1), the data were randomly split into 90% train and 10% test sets (2). The train data were pre-processed and the model hyperparameters were tuned using leave-one-out-cross-validation (3–4). Performance on the train data was calculated (5) and the set of hyperparameters with the highest kappa statistic were chosen for the final model (6). This model was then applied to the test data (7) and the test performance metrics were calculated (8). Steps 2-8 were repeated 100 times using bootstrap samples of the test and train sets, and the mean and 95% CI were calculated for test and train performance metrics (9). Scenario A: all titre predictor variables (pre- and post-infection PRNT titres and change in titre against all four dengue virus serotypes and Japanese encephalitis virus, and the number of days between measurement of the pre- and post-infection titres and the date of infection). Scenario B: all predictor variables (titre predictor variables plus age, year, and school). Scenario C: post-infection PRNT titre of the dengue serotypes were predictor variables. Scenario D: all titre predictor variables but only predicted seropositive cases. Seropositive individuals were defined as those with a pre-infection titre ≥ 10 for at least one DENV serotype. CI: confidence interval. PRNT: plaque reduction neutralisation test.

**Figure 5:**
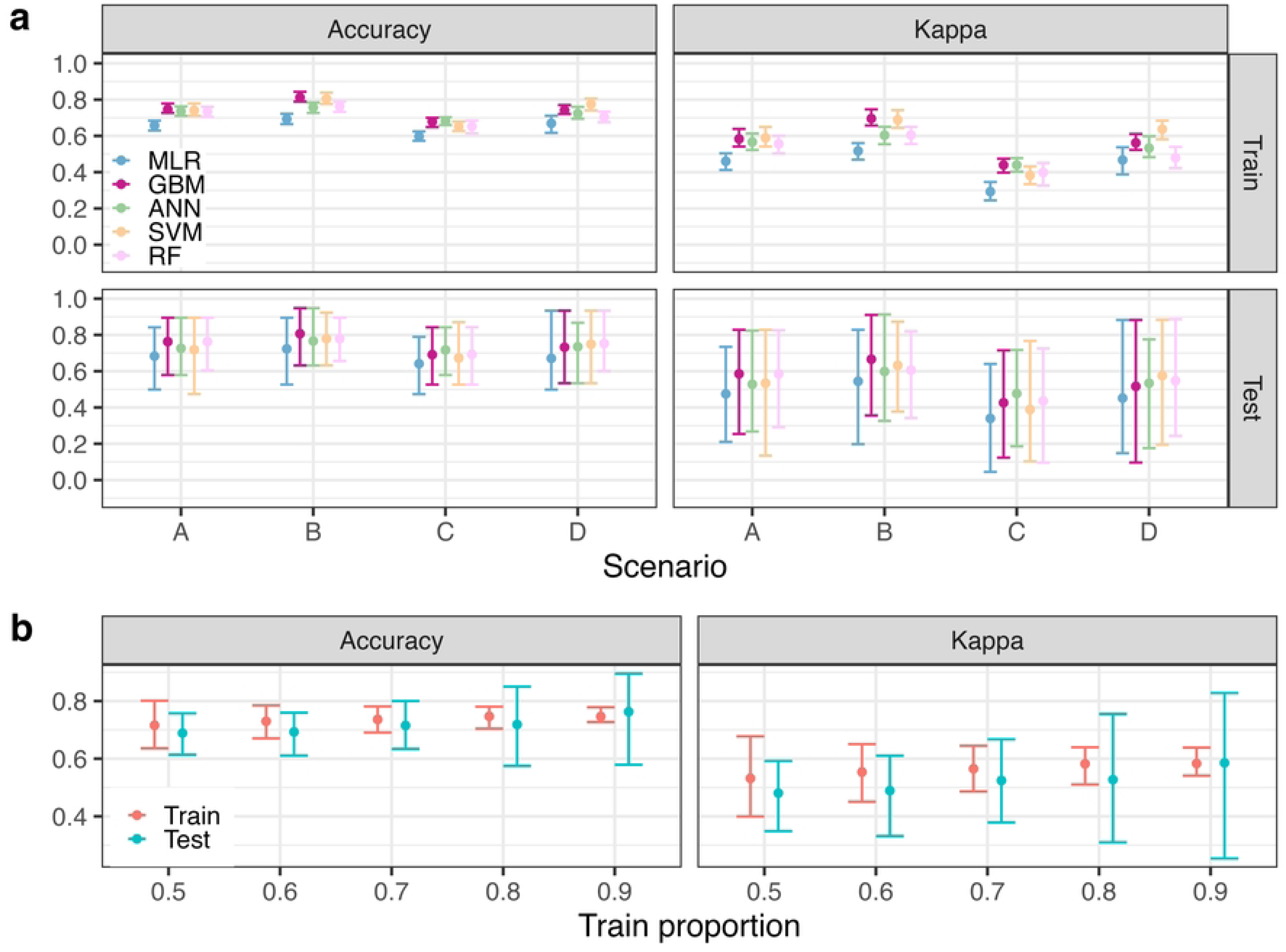
Comparison of regression and machine learning models for predicting the infecting DENV serotype. **a)** Performance of each model in predicting the infecting serotype across four scenarios using a 90/10 train/test split of the data. Scenario A and scenario D predictor variables were pre- and post-infection PRNT titres against DENV 1-4 and JEV, change in PRNT titre and number of days between infection and measurement of pre- and post-infection titres. Scenario B used the same predictor variables as A and C, plus year, age, and school. Scenario C predictor variables were post-infection PRNT against DENV 1-4. Scenario’s A, B, and C predicted all cases, scenario D predicted cases in seropositive individuals only. Seropositive individuals were defined as those with a pre-infection titre ≥ 10 for at least one DENV serotype. **b)** Comparison of different train/test proportions (90/10, 80/20, 70/30, 60/40, 50/50) for predicting the infecting dengue virus serotype (scenario A), using the GBM classification model. For each performance metric, the mean (bar) and 95% confidence interval (error bar) were calculated using 100 bootstrap samples of the test and train sets. Test performance was calculated by predicting on the hold-out-sample. Train performance was calculated using leave-one-out cross validation. ANN: artificial neural network. MLR: multinomial logistic regression. SVM: support vector machine. GBM: gradient boosting machine. RF: random forest. PRNT: plaque reduction neutralisation test. DENV: dengue virus.

In scenario B, we investigated whether the predictive performance could be further improved by including spatiotemporal variables in the model (i.e., the subjects age and school, and the year of infection). In this scenario, the best performing classifier was again GBM, with a test accuracy and kappa statistic of 80.6% (95% CI 63.2-94.7%) and 0.67 (95% CI 0.36-0.91), respectively (**Fig. 5a**). The train kappa statistic was 0.70 (95% CI 0.66-0.75), a significant increase compared to the train kappa statistics in scenario A. In scenario C, only the post-infection DENV PRNT titres were used as predictor variables (**Fig. 5a**). As expected, the overall test performance was lower than when using all predictors. The ANN classifier performed the best, with a test accuracy and kappa statistic of 71.7 % (95% CI 57.9- 84.2) and 0.48 (95% CI 0.19-0.72), respectively. Again, the machine learning classifiers outperformed MLR on the training and test sets, despite the large uncertainty in the performance estimates on the test set. Finally, the performance in scenario D was slightly lower than in scenario A and subject to greater uncertainty, reflecting the smaller dataset (N = 169) and prediction of only seropositive cases. Across all four scenarios, the GBM classifier performed the best.

**Table 1** shows the class performance of MLR and the machine learning models on the test set for predicting the infecting serotype in scenario A. Notably, the model sensitivity was higher for DENV-2 and DENV-3 than DENV-1, which is likely linked with the serotype-specific prevalence of infection observed in the test set, equal to 52.6%, 31.6%, and 15.8%, respectively. The SVM had the highest sensitivity against DENV-1 (65%), followed by MLR (61%). Conversely, the sensitivity of MLR against DENV-3 was notably lower than the machine learning models (49% vs. 54-66%). The highest positive predictive values (PPV) were achieved by GBM and ANN for DENV-1, SVM for DENV-2, and RF and GBM for DENV-3. The negative predictive values (NPV) were generally similar across classifiers, for each serotype (**Table 1**). The inclusion of the spatio-temporal variables (scenario B) increased the predictive performance against all serotypes. For instance, the sensitivity of GBM increased from 53%, 90%, and 66% to 58%, 93%, and 71% for DENV-1, DENV-2, and DENV-3, respectively (**S2 Table**). Therefore, it will be important to tailor the classification method and variables used to the specific setting.

**Table 1:**
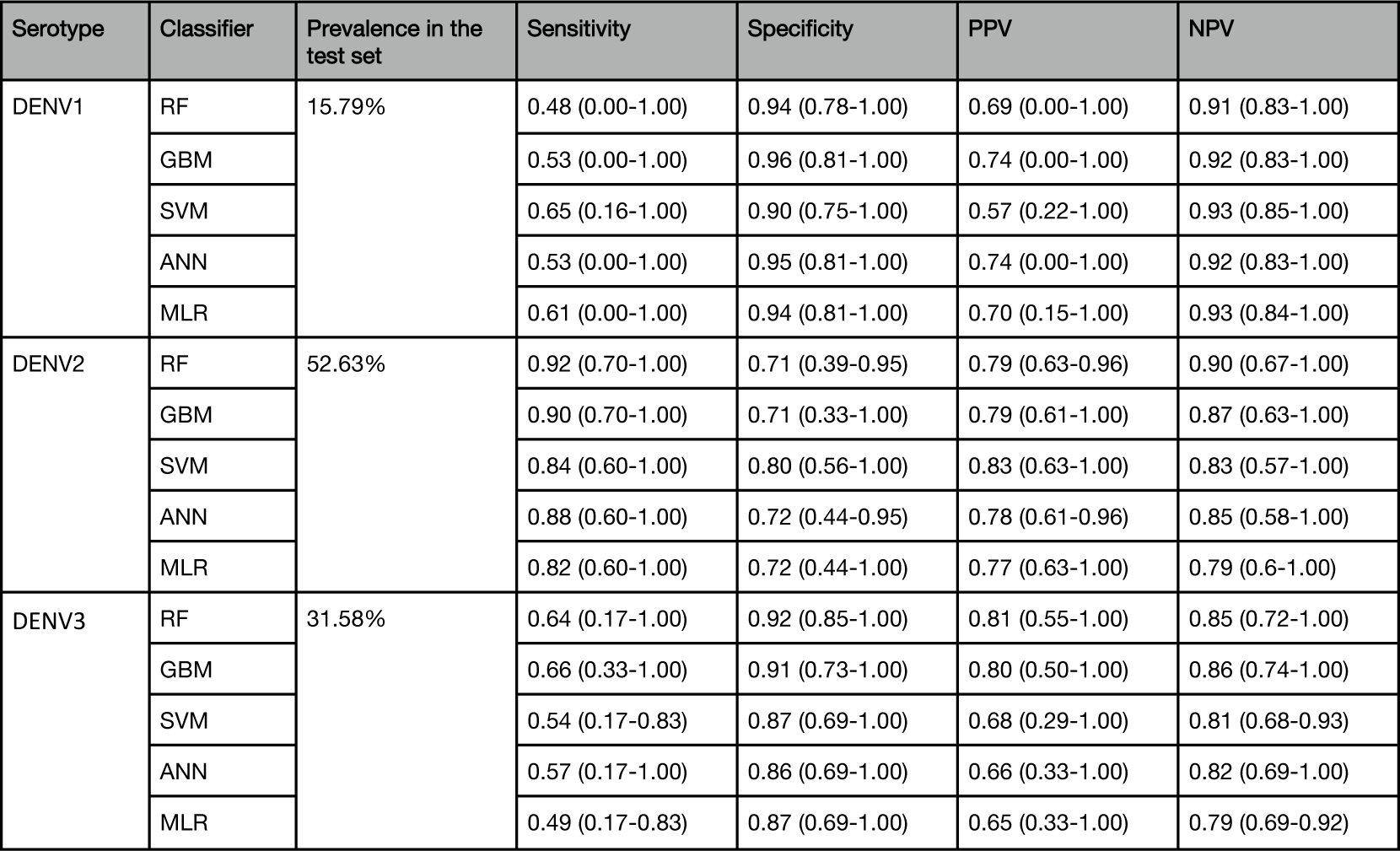
Mean (95% confidence interval) test class performance of regression and machine learning models for predicting the infecting DENV serotype using all titre predictor variables (Scenario A). For each performance metric, the mean and 95% confidence interval were calculated using 100 bootstrap samples of the hold-out-sample (10%). There were no DENV4 cases in the test set. ANN: artificial neural network. MLR: multinomial logistic regression. SVM: support vector machine. GBM: gradient boosting machine. RF: random forest. PRNT: plaque reduction neutralisation test. DENV: dengue virus. JEV: Japanese encephalitis virus. PPV: positive predictive value. NPV: negative predictive value.

| Serotype | Classifier | Prevalence in the test set | Sensitivity | Specificity | PPV | NPV |
| --- | --- | --- | --- | --- | --- | --- |
| DENV1 | RF | 15.79% | 0.48 (0.00-1.00) | 0.94 (0.78-1.00) | 0.69 (0.00-1.00) | 0.91 (0.83-1.00) |
|  | GBM |  | 0.53 (0.00-1.00) | 0.96 (0.81-1.00) | 0.74 (0.00-1.00) | 0.92 (0.83-1.00) |
|  | SVM |  | 0.65 (0.16-1.00) | 0.90 (0.75-1.00) | 0.57 (0.22-1.00) | 0.93 (0.85-1.00) |
|  | ANN |  | 0.53 (0.00-1.00) | 0.95 (0.81-1.00) | 0.74 (0.00-1.00) | 0.92 (0.83-1.00) |
|  | MLR |  | 0.61 (0.00-1.00) | 0.94 (0.81-1.00) | 0.70 (0.15-1.00) | 0.93 (0.84-1.00) |
| DENV2 | RF | 52.63% | 0.92 (0.70-1.00) | 0.71 (0.39-0.95) | 0.79 (0.63-0.96) | 0.90 (0.67-1.00) |
|  | GBM |  | 0.90 (0.70-1.00) | 0.71 (0.33-1.00) | 0.79 (0.61-1.00) | 0.87 (0.63-1.00) |
|  | SVM |  | 0.84 (0.60-1.00) | 0.80 (0.56-1.00) | 0.83 (0.63-1.00) | 0.83 (0.57-1.00) |
|  | ANN |  | 0.88 (0.60-1.00) | 0.72 (0.44-0.95) | 0.78 (0.61-0.96) | 0.85 (0.58-1.00) |
|  | MLR |  | 0.82 (0.60-1.00) | 0.72 (0.44-1.00) | 0.77 (0.63-1.00) | 0.79 (0.6-1.00) |
| DENV3 | RF | 31.58% | 0.64 (0.17-1.00) | 0.92 (0.85-1.00) | 0.81 (0.55-1.00) | 0.85 (0.72-1.00) |
|  | GBM |  | 0.66 (0.33-1.00) | 0.91 (0.73-1.00) | 0.80 (0.50-1.00) | 0.86 (0.74-1.00) |
|  | SVM |  | 0.54 (0.17-0.83) | 0.87 (0.69-1.00) | 0.68 (0.29-1.00) | 0.81 (0.68-0.93) |
|  | ANN |  | 0.57 (0.17-1.00) | 0.86 (0.69-1.00) | 0.66 (0.33-1.00) | 0.82 (0.69-1.00) |
|  | MLR |  | 0.49 (0.17-0.83) | 0.87 (0.69-1.00) | 0.65 (0.33-1.00) | 0.79 (0.69-0.92) |

To further understand the class imbalance, **Fig. 6** presents the predicted and true infecting serotype in cases misclassified by the GBM model in scenario A. The most frequent misclassification combination was a DENV-3 infection predicted to be DENV-2, which may be because the greatest change in PRNT titre following a DENV-3 infection was equally likely to be DENV-2 as DENV-3 (**Fig.2**). 11.1% of misclassifications occurred in cases where the highest post-infection titre was tied between the predicted and true infecting serotype. Of the remaining misclassifications, the titre of the infecting serotype was higher than the predicted serotype in 65.4% of the misclassified cases (**Fig. 6**).

**Figure 6:**
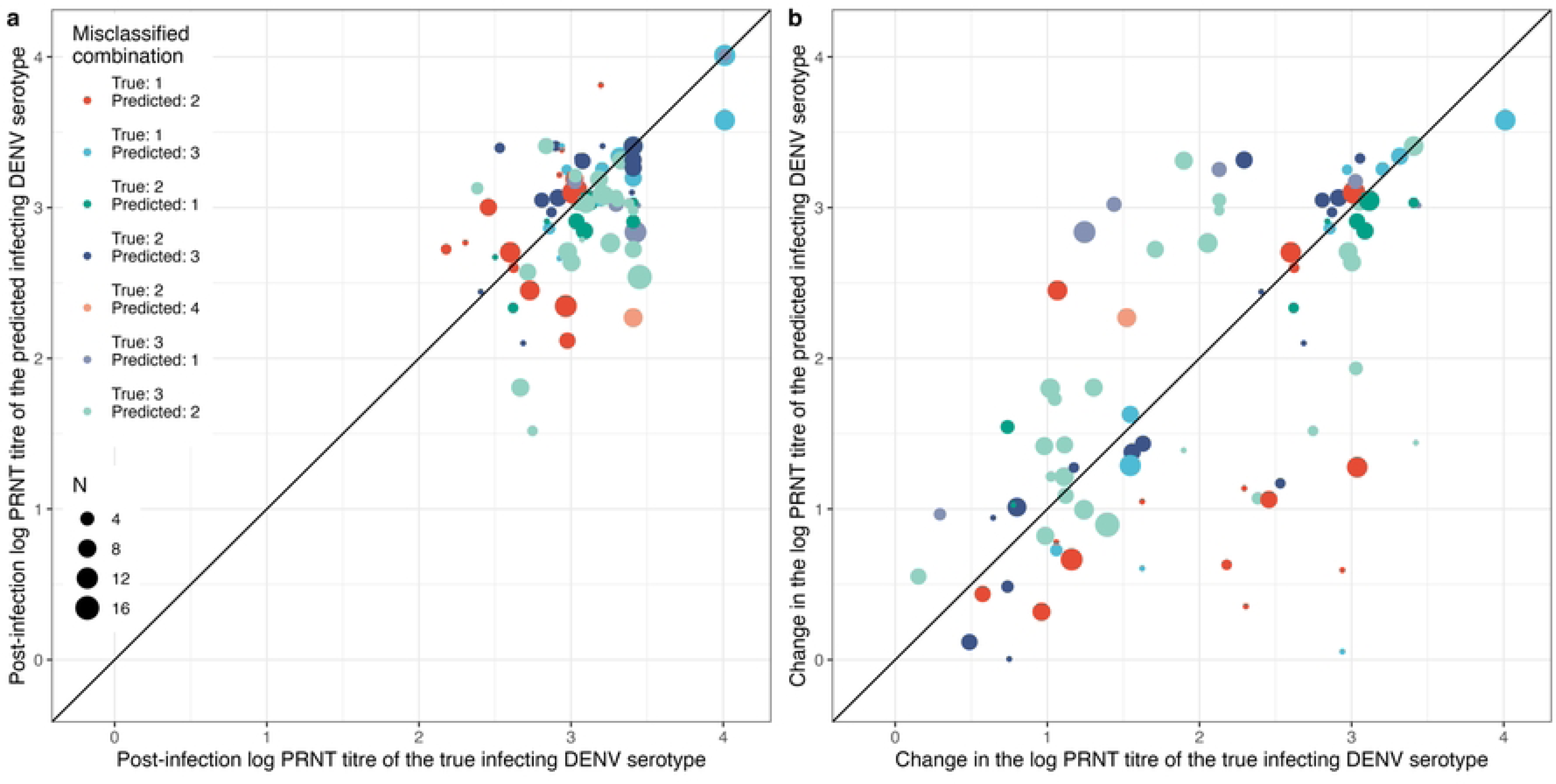
Post-infection (a) and change (b) in log PRNT titres of the predicted infecting DENV serotype (y axis) and the true infecting DENV serotype (x axis) in cases that were misclassified by the gradient boosting machine model. N = the number of times a case was misclassified out of 100 bootstrap samples of the test set. Predictor variables were pre- and post-infection PRNT titres against DENV 1-4 and JEV, change in PRNT titre and number of days between infection and measurement of pre- and post-infection titres (scenario A). PRNT: plaque reduction neutralisation test. DENV: dengue virus. JEV: Japanese encephalitis virus.

### Sensitivity analysis

In a sensitivity analysis, we set any titre value above 2560 to 2560, to investigate a discrepancy in the maximum PRNT dilution. Although the distribution of titres was similar to the main analysis (**Fig. 1, and S2b Fig.**), 47 seropositive cases had tied highest titre values, compared to 29 in the main analysis (**Fig. 2a, S2a Fig.**). Notably, this led to a reduction in classification performance, with a test kappa statistic of 0.47 (95% CI 0.12-0.78) in scenario A (**S2c Fig.**).

## Discussion

Despite the development of multiple dengue vaccines, the lack of standardised interpretation of neutralising antibodies still hinders the identification of correlates of protection. Whilst the primary endpoint of dengue vaccine trials has so far focussed on virologically confirmed disease or hospitalisation, the availability of methods that can identify subclinical infections can pave the way to the assessment of serotype-specific protection against infection. Here, we trained machine learning classification models to predict the infecting DENV serotype from pre- and post-infection neutralising antibody titres, achieving a mean accuracy of 76.3% using the GBM classifier. When additionally including information on the individual’s school, age and year of infection, the accuracy increased to 80.6% on average. Consistent with original antigenic sin (6), we found that 57.6% of the highest post-infection titres of seropositive cases correspond with the serotype of the highest pre-infection titre, rather than to the infecting serotype. Despite this, a classification model trained only on post-infection DENV titres was able to predict the infecting serotype with 71.7% accuracy, on average.

Very few of the cases in seronegative individuals exhibited a monotypic response (20,22,23), implying that a higher specificity (e.g., PRNT-70 or -90) may be needed to disentangle cross-reactive antibodies, even in primary cases. However, the highest post-infection antibody titres of seronegative individuals were frequently against a non-infecting serotype. This may be due to the antigenic overlap between serotypes, for instance the highest titre following multiple DENV-1 infections was DENV-3 and vice-versa, which are phylogenetically the most similar (24). However, previous work suggests that the highest neutralising response following infection in seronegative individuals is against the infecting serotype (6). An alternative explanation is that the seronegative individuals had prior dengue exposure, but their antibody levels decayed below the limit of detection, causing their misclassification as seronegative. Alternatively, it is possible that the DENV strain used in the PRNT differed from the circulating strain.

The classification models developed in this study can be extended to investigate the association between sequences of serotype-specific infections and disease protection or enhancement (25). Furthermore, these models could be used to identify subclinical infections within vaccine trials using serology data, allowing the estimation of serotype-specific vaccine efficacy against infection which has the potential to alter serotype transmission dynamics (26). Additionally, they could be used to investigate the prevalence of homotypic reinfection, which may be more frequent than previously realised, especially if asymptomatic (3,27,28).

We found that inclusion of the infection year, participant age, and school as predictor variables, in addition to PRNT titres and the duration of time between infection and titre measurement, increased the predictive performance of the classification models. As these predictor variables are specific to the current cohort, their inclusion in the model and the use of a single dataset reduces the generalisability, highlighting the need for more individual-level serological data from epidemiological studies and vaccine trials.

This study builds on previous work by Van Panhuis *et al.* (21) who used MLR to infer the infecting serotype from neutralising antibody titre data. We found that machine learning classifiers perform better on average than MLR, reflecting their ability to model complex, non-linear relationships. However, this comes with the cost of reduced model simplicity and interpretability. Additionally, there was wide variability of the test set performance, attributable to the small sample size of the dataset. Machine learning is expected to outperform MLR on larger datasets, owing to its flexibility in describing complex relationships, but this analysis also shows that to draw definitive conclusions, machine learning algorithms require more data. MLR and the classifiers showed variable performance against each serotype, reflecting the class imbalance of the data. Future work predicting the infecting serotype from larger or multiple data sets with more equal representation of each serotype is likely to improve overall performance, as well as the performance against each serotype.

A further limitation is the inconsistent maximum dilutions used for the PRNT. We addressed this in a sensitivity analysis by setting any titre values >2560 to 2560, which resulted in a loss of predictive performance. To capture the full range of antibody dynamics, especially in seropositive infections, serial dilutions should therefore be performed to at least 1:10240 or criteria pre-specified, dictating the sequence with which different dilutions should be completed. Finally, the date between measurement of titres and the date of infection ranged from 9-281 days. Although we included this as a predictor variable, increasing performance, it is possible that subclinical infections or boosting of titres may have occurred within that time, further complicating interpretation of PRNT titres.

This analysis compared the performance of four machine learning classifiers and MLR in predicting the infecting serotype from neutralising antibody titres. Machine learning classifiers predicted the infecting serotype with high accuracy, overcoming challenges in interpreting cross-reactive PRNT titres, in both seropositive and seronegative cases. If applied to longitudinal antibody titre data, the models developed here can be used to predict sequential infections, subclinical infections, and homotypic reinfections, making them important tools for our understanding of dengue immunity, disease dynamics and help identify serotype-specific immune correlates of vaccine induced protection.

## Methods

### Data Collection

Antibody response data were obtained from a prospective cohort study conducted on children in grades 1-5 in Kamphaeng Phet, Thailand, as previously described (29). Briefly, the study enrolled children from 12 schools in 1998, with new first graders enrolled each January until 2002. Active case surveillance for acute dengue illness ran from June to November, during which three blood samples (June, August, and November) were collected for dengue serology. Village healthcare workers visited absent students to check for fever within seven days or a temperature >38°C, and a referral to the public health clinic was made, if necessary, where an acute illness blood sample was obtained. Children who reported directly to the public health clinic or were admitted to the hospital were also evaluated for acute dengue illness.

Acute dengue infection was confirmed through dengue serology (immunoassays against DENV-1-4 and Japanese encephalitis virus (JEV)) or detection of the virus (29). Confirmation of the infecting serotype was performed through RT-PCR within six days of acute DENV illness. PRNT titres were measured from pre- and post-infection blood samples of children with confirmed dengue infection. All PRNT titres were performed in the same lab using the same protocol (29). A monolayer of LLC-MK2 cells were infected with a constant amount of DENV, in the presence of four-fold serial dilutions of heat-inactivated plasma from patients (1:10-1:2560 or 1:10-1:10240). The viral strains used in the PRNT were DENV-1 (16007), DENV-2 (16681), DENV-3 (16562), and DENV-4 (1036), and JEV (Vaccine strain; SA 14-14-2). PRNT_50_ were calculated using probit regression and reported as the reciprocal titre.

Informed assent and consent were obtained from all participants or guardians, and all data used in this present study was de-identified.

### Model development and validation

We compared the performance of four non-parametric classifiers (RF, GBM, ANN, and SVM) to MLR in predicting the infecting DENV serotype from neutralising antibody titres.

RF and GBM are ensemble decision trees, hierarchical structures which break down the classification process into a series of sequential conditions. RF builds many trees from bootstrap training data samples, so each tree is grown on random subsets of the data, with a random subset of variables considered at each decision node. Each tree provides a class vote to classify observations. The class with the majority votes is selected for the final prediction. By combining the predictions of many de-correlated trees, RF reduces variance and the risk of overfitting, improving predictive performance. Whereas RF builds many trees simultaneously, GBM builds decision trees sequentially. Starting with a weak learner (i.e., a tree with only a few splits), GBM aims to increase the performance with each new tree, by fitting to observations misclassified by the previous trees. The final model is a linear combination of many trees, that provide a class vote, like RF. GBM can reduce both the bias and variance of single decision trees; however, they may require more data than RF.

Support vector classifiers create linear boundaries, or hyperplanes, to divide the data into different classes. The optimal hyperplane is found by optimising the margin between the plane and the nearest training observations. SVM extends this, by using kernels to handle non-linear decision boundaries, which enlarge the feature space into many dimensions. This makes SVM effective in high dimensional space, although they can underperform with noisy data. A one-versus-one approach is used to classify more than two classes.

ANNs are inspired by the neural networks in the brain and compromise of node layers, which are analogous to neurons. The output layer (the classification) is modelled from the input layer (the predictor variables) via an intermediary layer of hidden nodes. In this analysis, a single hidden layer neural network was implemented. The hidden layer derives non-linear transformations from the input layer. Weight decay is used to regularise ANN models, which are otherwise prone to over-fit due to the high number of estimated parameters (intercepts and weights). Fitting multiple independent models and aggregating their predictions also reduces variance.

The outcome variable was the infecting serotype (DENV-1-4), identified through RT-PCR. The titre predictor variables were the pre- and post-infection PRNT titres, change in titre and time between infection and measurement of titres (**Supplementary Table 1**). Additionally, the year of infection and the age and school of the participant were considered as predictor variables. Titres recorded as “<10”, “>2560” or “>10240” were set to 0, 2560 and 10240, respectively. All PRNT titres were log_10_ transformed after adding 1, to obtain a normal distribution. For SVM and ANN, the log-transformed titres were centred and scaled.

**Figure 1** presents an overview of model development and validation. For each classifier, four modelling scenarios were investigated. Scenario A used all titre measurements as predictor variables and predicted all cases, scenario B used all titre measurements plus year of infection and the age, and school of the individuals and predicted all cases, scenario C used post-infection DENV titres as predictor variables and predicted all cases, and scenario D used all titre measurements as predictor variables and predicted only seropositive cases (pre-infection titre ≥ 10 to at least one serotype) (**Supplementary Table 1**). We did not consider feature selection due to the small number of predictor variables.

Model performance was evaluated on out-of-sample datasets by splitting the dataset, so the model was trained on 90% of the data (train set) and validated on 10% of the data (test set), while preserving the overall serotype distribution within both sets. The training set was used to tune the hyperparameters of each classifier using leave-one-out-cross-validation. (**Supplementary Table 2**). The hyperparameter combination that gave the best performance, measured using the kappa statistic, was chosen to predict the infecting serotypes of the test set. In total, 100 estimates of the train and test performance were obtained using 100 bootstrap samples of the test and train sets, from which we calculated the mean and 95% CI (2.5-97.5 percentiles). The analysis was conducted through the caret package (30), using R studio version 1.3.1093 (31). The packages for each classifier are nnet(32), kernlab (33), gbm (34), ranger (35).

### Sensitivity analyses

A sensitivity analysis was conducted to ascertain the impact of different train and test splits on model performance and uncertainty. To this end, the GBM algorithm was applied to scenario A using the following test and train splits: 10%/90% (baseline), 20%/80%, 30%/70%, 40%/60%, 50%/50%.

Finally, the maximum dilution for the PRNT of some antibody samples was 1:2560, whilst for others it was 1:10240. No samples collected following a DENV-2 infection were diluted above 1:2560. To investigate whether the different dilutions impacted the classification performance, we conducted a sensitivity analysis where we set any sample greater than 2560 to 2560.

## Contributors

BCD contributed to the conceptualisation, formal analysis, methodology, visualisation, writing – original draft and revisions. DB contributed to data curation, investigation, writing – review & editing. TH contributed to data curation, investigation, writing – review & editing. SI contributed to data curation, investigation, writing – review & editing. AF contributed to the funding acquisition, investigation, project administration, writing -revisions. DATC contributed to the methodology, writing – review & editing. KA contributed to the funding acquisition, investigation, project administration, methodology, writing – review & editing. ID contributed to the conceptualisation, methodology, writing – original draft and revisions.

## Data sharing

Data and code are provided to fit the models using all titre data (scenario A) and using all titre data plus year of infection and age-group (modified scenario B). School and age in years are not published to ensure anonymity. Data and code are available at: https://github.com/bnc19/dengue-serotype-classification-public

## Acknowledgments

ID acknowledges funding by the Wellcome Trust (grant number 213494/Z/18/Z). BCD acknowledges PhD funding from the UK Medical Research Council. ID and BCD acknowledge funding from the MRC Centre for Global Infectious Disease Analysis (reference MR/X020258/1), funded by the UK Medical Research Council (MRC). This UK funded award is carried out in the frame of the Global Health EDCTP3 Joint Undertaking. The original cohort study was funded by NIH grant P01 A1034533 and the Military Infectious Disease Research Program. KA was supported by a grant from the US NIH 1R01AI175941. DATC was supported by a grant from the US NIH R01AI175495.

Material has been reviewed by the Walter Reed Army Institute of Research. There is no objection to its presentation and/or publication. The opinions or assertions contained herein are the private views of the author, and are not to be construed as official, or as reflecting true views of the Department of the Army or the Department of Defense. The investigators have adhered to the policies for protection of human subjects as prescribed in AR 70–25.

